# COmplexome Profiling ALignment (COPAL) reveals remodeling of mitochondrial protein complexes in Barth syndrome

**DOI:** 10.1101/411660

**Authors:** Joeri Van Strien, Sergio Guerrero-Castillo, Iliana A. Chatzispyrou, Riekelt H. Houtkooper, Ulrich Brandt, Martijn A. Huynen

## Abstract

**Motivation:** Complexome profiling combines native gel electrophoresis with mass spectrometry to obtain the inventory, composition and abundance of multiprotein assemblies in an organelle. Applying complexome profiling to determine the effect of a mutation on protein complexes requires separating technical and biological variations from the variations caused by that mutation.

**Results:** We have developed the COmplexome Profiling ALignment (COPAL) tool that aligns multiple complexome profiles with each other. It includes the abundance profiles of all proteins on two gels, using a multidimensional implementation of the dynamic time warping algorithm to align the gels. Subsequent progressive alignment allows us to align multiple profiles with each other. We tested COPAL on complexome profiles from control mitochondria and from Barth syndrome (BTHS) mitochondria, which have a mutation in tafazzin gene that is involved in remodelling the inner mitochondrial membrane phospholipid cardiolipin. By comparing the variation between BTHS mitochondria and controls with the variation among either, we assessed the effects of BTHS on the abundance profiles of individual proteins. Combining those profiles with gene set enrichment analysis allows detecting significantly affected protein complexes. Most of the significantly affected protein complexes are located in the inner mitochondrial membrane (MICOS, prohibitins), or are attached to it (the large ribosomal subunit).

**Availability and implementation:** COPAL is written in Python and is available from gttp://github.com/cmbi/copal.

## INTRODUCTION

Complexome profiling combines native gel electrophoresis with protein mass spectrometry to obtain a comprehensive inventory of the proteins and multiprotein complexes (the complexome) in a biological sample. First, blue-native gel electrophoresis separates intact protein complexes from each other (Ferrandez et al., 2012; Schägger and von Jagow, 1991). After cutting the gel lanes into slices, these are subjected to tryptic digest and mass spectrometry, yielding an abundance profile reflecting the migration pattern in the native gel per each identified protein. Similar abundance profiles indicate interaction between proteins. Agglomerative clustering based on profile-similarity can thus be used to assemble protein complexes from the dataset (Giese et al., 2015; Heide et al., 2012).

Complexome profiling is rapidly becoming a versatile research tool that among others has been employed to discover new complexes (König et al., 2016; Wai et al., 2016), to identify new proteins in known complexes (Huynen et al., 2016; Sanchez-Caballero et al., 2016), to study the assembly of protein complexes (Guerrero-Castillo et al., 2017a; Vidoni et al., 2017), and to detect tissue specific splice variants in protein complexes (Guerrero-Castillo et al., 2017b). Complexome profiling has initially been developed for mammalian mitochondria, but has been shown to be applicable also to other cellular systems, like the V-type H^+^-ATPase (Van Damme et al., 2017), the endoplasmatic reticulum (Prior et al., 2016), plant mitochondria (Senkler et al., 2017) and bacteria (de Almeida et al., 2016; Schimo et al., 2017).

Here we introduce a bioinformatics tool to detect how disease-causing mutations alter the mitochondrial complexome. Given the wealth of data contained in complexome profiles, a comprehensive comparison of control complexome profiles with the complexome profiles of cells carrying a mutation requires standardized, computational procedures to prioritize the proteins and protein complexes most affected by a given mutation. Furthermore, it requires tools to separate technical and biological variations from the ones caused by the mutation. One technical variation is B an overall shift in the migration patterns between different complexome profiling gels. Furthermore the number of slices the gels are cut into varies between labs, ranging from 56 (Wohlbrand et al., 2016) or 66 (de Almeida et al., 2016) to 70 (Senkler et al., 2017) or even up to 230 (Müller et al., 2016), complicating large-scale comparisons. Aligning the gels is particularly important when comparing complexome profiles from cells harboring a mutation with control complexome profiles, because mutations can cause changes in the composition of protein complexes and thus shifts in their migration pattern. One would like to separate such shifts, affecting a limited number of protein complexes, from technical variations that affect the overall migration pattern. To this end, we have developed a COmplexome Profiling ALignment (COPAL) tool that uses a multidimensional implementation of dynamic time warping (Sakoe and Chiba, 1978; ten Holt et al., 2007) to combine the signals of all proteins and effectively align the gels *in silico*. By employing a progressive alignment technique, we subsequently align multiple lanes from different gels with each other.

Based on this “multiple gel alignment” we can automatically quantify the differences per protein between patients’ cells and control cells and thus prioritize the proteins most affected by a mutation. Furthermore, COPAL allows us to use an integrative technique like Gene Set Enrichment Analysis (Subramanian et al., 2005) to prioritize the protein complexes that are significantly affected by mutations.

We tested our approach on mitochondrial complexome profiles of four patients with Barth syndrome (Houtkooper et al., 2009). Barth syndrome (BTHS) is associated with mitochondrial cardiomyopathy and is characterized at the molecular level by defective cardiolipin (CL) remodeling, a phospholipid that makes up up to 20% of the inner mitochondrial membrane. It is a particularly interesting disease to study with complexome profiling because of the many reported interactions of CL with mitochondrial proteins and protein complexes of the inner mitochondrial membrane (Paradies et al., 2014). Our approach allowed us to align the profiles of mitochondria of fibroblasts of the patients with those of five controls, four of which were run on different gels, and to automatically detect the proteins and protein complexes significantly affected by the mutations.

## MATERIAL AND METHODS

### Complexome Profiling Datasets

Human fibroblasts were cultured and mitochondria were isolated as described previously (Huynen et al., 2016). ESI-MS/MS mass spectrometry and acquisition of complexome profiles was performed as described in (Guerrero-Castillo et al., 2017a). Complexome datasets of four patients carrying mutations in the Tafazzin gene (Table S1) (Houtkooper et al., 2009) and two of the healthy controls (C1_G1 and C1_G3) were taken from (Chatzispyrou et al., 2018) and from (Chatzispyrou et al., 2017) respectively. The unprocessed complexome profiles of the three other healthy controls (C1_G2, C2_G2, C3_G2) and of C1_G3 are provided as supplementary data (Table S2). C1-C3 denotes the control fibroblast and G1-G3 the blue-native gel electrophoresis run from which the lane for the complexome profile was taken.

### Normalization

The complexome profile data of a single sample is a two dimensional matrix with the intensity based absolute quantification (iBAQ) (Schwanhausser et al., 2011) values per protein (first dimension) and per gel slice (second dimension). In order to correct for varying overall intensities between the gels, iBAQ values are normalized per sample such that the summed up iBAQ values of the detected proteins over all the samples are the same. For the analysis described here, we used an enriched mitochondrial preparation. Normalization and overall analysis were limited to mitochondrial proteins from MitoCarta2.0 (Calvo et al., 2016) to prevent varying, non-mitochondrial contaminants affecting the results.

### Complexome Profile Alignment

We developed COmplexome Profiling Alignment (COPAL) to align the gels using the normalized iBAQ values of all mitochondrial proteins. COPAL is based on dynamic time warping (Sakoe and Chiba, 1978). Classically, dynamic time warping aligns two series of values, e.g. two time series, by finding in the matrix the “warp path” with the lowest total distance, also known as the global cost. In dynamic time warping, the local distance between two points is the difference between the corresponding values in the time series. In our COPAL implementation of dynamic time warping, the local distance between two profiles at a certain point in the matrix is the sum, for *all* proteins, of the differences between the iBAQ values per protein (Figure 1). Such time warping has been named “multi-dimensional time warping” (ten Holt et al., 2007). Note that proteins with high iBAQ values, which contain the strongest signal, contribute most to this sum of differences and therewith to the alignment. After determining the locally optimal warp path at each location, the dynamic time warping algorithm traces back along the path resulting in the lowest global cost, providing the optimal alignment between two samples. The global cost is a measure for the total difference between two profiles and does not include gap penalties. It is therewith independent of technical variations in the overall migration patterns between gels and can be used to quantify the difference between complexome profiles.

**Figure 1.**
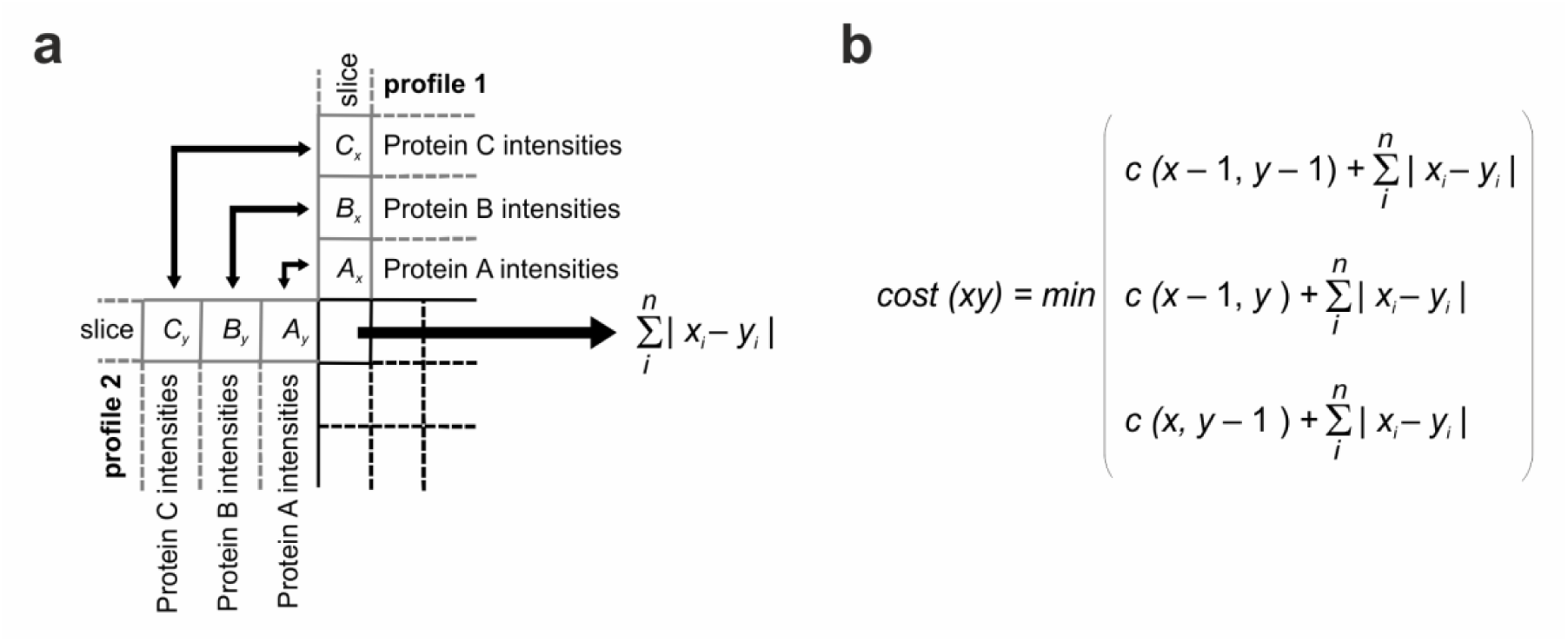
COPAL procedure for the alignment of two complexome profiling samples. A) The distance, or “local cost” between two gel slices is the sum of the absolute differences of the iBAQ values for all proteins on that slice. Here, *i* is an index for each protein, and *n* is the total number of proteins in the complexome profile, x and y are intensities of proteins from profile 1 and profile 2. B) As is the case in classical time warping, the cost(xy) is the minimum of the local cost at point (x,y) plus one of three preceding costs: the one above it (x, y-1) corresponding to a slice (gap) inserted in the alignment of gel y, the one beside it (x-1,y), corresponding to a slice inserted in gel x, or the one diagonal from it (x-1, y-1) corresponding to no slice insertion. After building up the matrix in this manner the two gels are aligned by tracing back along the optimal warping path, as in standard dynamic time warping, thus obtaining the alignment with the minimal global cost.

## Progressive alignment

To align more than two complexome profiles with each other we used a progressive alignment approach (Hogeweg and Hesper, 1984) similar to the one used by clustalW (Thompson et al., 1994). The global cost values from pairwise alignments between all profiles are used to create a guide tree that determines the progressive alignment order. To align two alignments to each other, local distances between alignments need to be determined. The local distance between two alignments at a certain point is the average local distance of all sample pairs between the two alignments. After the final alignment is reached, the resulting sample-specific gap locations are used to “warp” each sample profile. At gap locations (inserted slices), the iBAQ value of every protein at the slice preceding the gap in that sample is repeated.

### Quantifying differences between abundance profiles per protein

We use the Hausdorff distance (Hausdorff, 1914) to quantify the difference between aligned abundance profiles per protein. Given two curves, or sets of coordinates, the Hausdorff distance is defined as the largest Euclidian distance from one set to the closest point in the other set. It thus “finds” the area where two abundance profiles differ the most, and measures this difference. It is a robust method for finding differences between point sets, like time series or, in this case, migration patterns (Moeckel and Murray, 1997).

As the Hausdorff distance is a Euclidean distance, it considers differences on both axes equally: i.e. it gives equal weight to shifts in the migration pattern of a protein corresponding to the protein complex mass, measured along the x-axis, as to shifts in its abundance, measured along the y-axis. Nevertheless, in the 2D plane of migration patterns, the (normalized) iBAQ values are many orders of magnitude higher than the length of the x-axis that is maximally the number of slices in which the gel is cut plus the number of inserted slices (gaps). To increase the contribution of shifts in migration patterns to the Hausdorff distance, we normalize the data per protein so that the maximum intensity value per protein over all the gels compared is equal to the length of the final alignment, creating a square plane in which the Hausdorff distance is calculated. The scaling parameter that is in this normalization can in principle also be used to weigh shifts in mass higher than shifts in intensity, and can be adapted in COPAL. In the analysis presented below we also normalized the abundance data to have a maximum ¼ as high as the final alignment length, effectively weighing changes in protein complex mass four-fold higher than changes in protein abundance. While the results per protein differed slightly, the set of significantly affected protein complexes remained the same.

### Calculating the difference between control and mutant complexomes per protein

To select for proteins with the largest difference in migration between *TAZ* mutation fibroblast mitochondria, hereafter names BTHS mitochondria, and controls, while showing consistent migration patterns within either group, we calculated the effect of the mutation by taking the average Hausdorff distance between the two groups and dividing them by the average Hausdorff distance within the groups (Eq1, where *B* and *W* are the number of protein sample pairs between and within groups, respectively). Proteins were subsequently sorted based on this score.

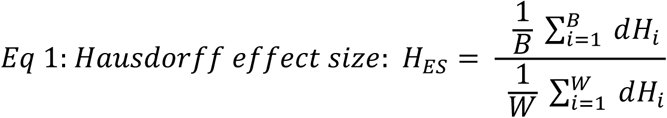

### COPAL

COPAL is written in python (version 2.7.6) and uses the pandas data analysis toolkit (version 0.20.1). It has a graphical user interface that is built using TkInter, the native python GUI package. The standalone executable tool is packaged with pyinstaller (version 3.2.1). Copal requires complexome profiles in excel or csv text document format as input. It outputs aligned complexome profiles as formatted excel files, as well as text files. Optionally, the tool also computes Hausdorff effect sizes between predefined controls and test complexomes. Extensive documentation of the program is included with the program itself at gttp://github.com/cmbi/copal.

### Gene set Enrichment Analysis to determine significantly affected protein complexes

To prioritize and integrate differences between complexome profiles at the level of protein complexes, we used gene set enrichment analysis (GSEA) (Subramanian et al., 2005) on the list of proteins that were rank ordered based on the Hausdorff effect size (Eq1). Given the direct interaction of CL with mitochondrial protein complexes we used the CORUM comprehensive resource of mammalian protein complexes (Ruepp et al., 2010), with manual corrections based on more recent literature (Supplementary methods), to examine enriched gene sets.

## RESULTS

### Complexome profiling of mitochondria from fibroblasts with and without *TAZ* mutations

We compared four complexome profiles from BTHS mitochondria carrying different mutations in the *TAZ* gene (Table S1) with five profiles from healthy controls, see (Chatzispyrou et al., 2018) for the four BTHS mitochondria complexomes and one control; Table S2 for the other four controls). Complexome profiles of all four patients and of one batch of mitochondria from control fibroblasts (C1_G1) were obtained from lanes of the same blue-native electrophoresis (BNE) gel (G1). Additional independently acquired complexome profiles of control C1 run in two separate gels (C1_G2, C1_G3) as well as two other control fibroblasts run on the second gel (C2_G2 and C3_G2) were included. Migration profiles were determined by in-gel tryptic digest and mass spectrometric analysis of 60 slices per lane. In total 4342 proteins were identified, of which 894 proteins (including proteins from alternative transcripts) were derived from 845 genes listed in MitoCarta 2.0 (Calvo et al., 2016). Of genes with multiple proteins the protein with the highest expression in the controls and BTHS mitochondria was retained for the analysis.

Relative protein abundances in each slice expressed as iBAQ values were determined by label-free quantification (Schwanhausser et al., 2011).

### Aligning complexome profiles

To make the relative protein abundance values comparable across the complexome profiles, we normalized the individual iBAQ values using a correction factor deduced from the sum of iBAQ values for all detected mitochondrial proteins in each profile. Subsequently we performed pairwise alignments of all profiles using COPAL. As expected, clustering the complexomes based on their alignment score showed that the largest differences occurred between the different BNE gels (Figure 2a). Progressive alignment based on this clustering resulted in a final alignment with four insertions per gel, distributed over thirteen different locations (Figure 2b, e, Table S3). Profiles that were run on the same gel tend to have slices inserted at the same locations, reflecting overall shifts between gels (Figure 2c, d) rather than within gels. The summed-up profiles of all mitochondrial proteins before and after alignment and normalization by COPAL, represented as abundance plots (bottom) or as heat maps (top) show how the procedure increases overall similarity between the same fibroblast mitochondria run on three different gels. The profiles of two mitochondrial proteins not affected by the mutations show how the procedure that is based on all proteins, corrected the relative position of two individual proteins (Figure S1).

**Figure 2.**
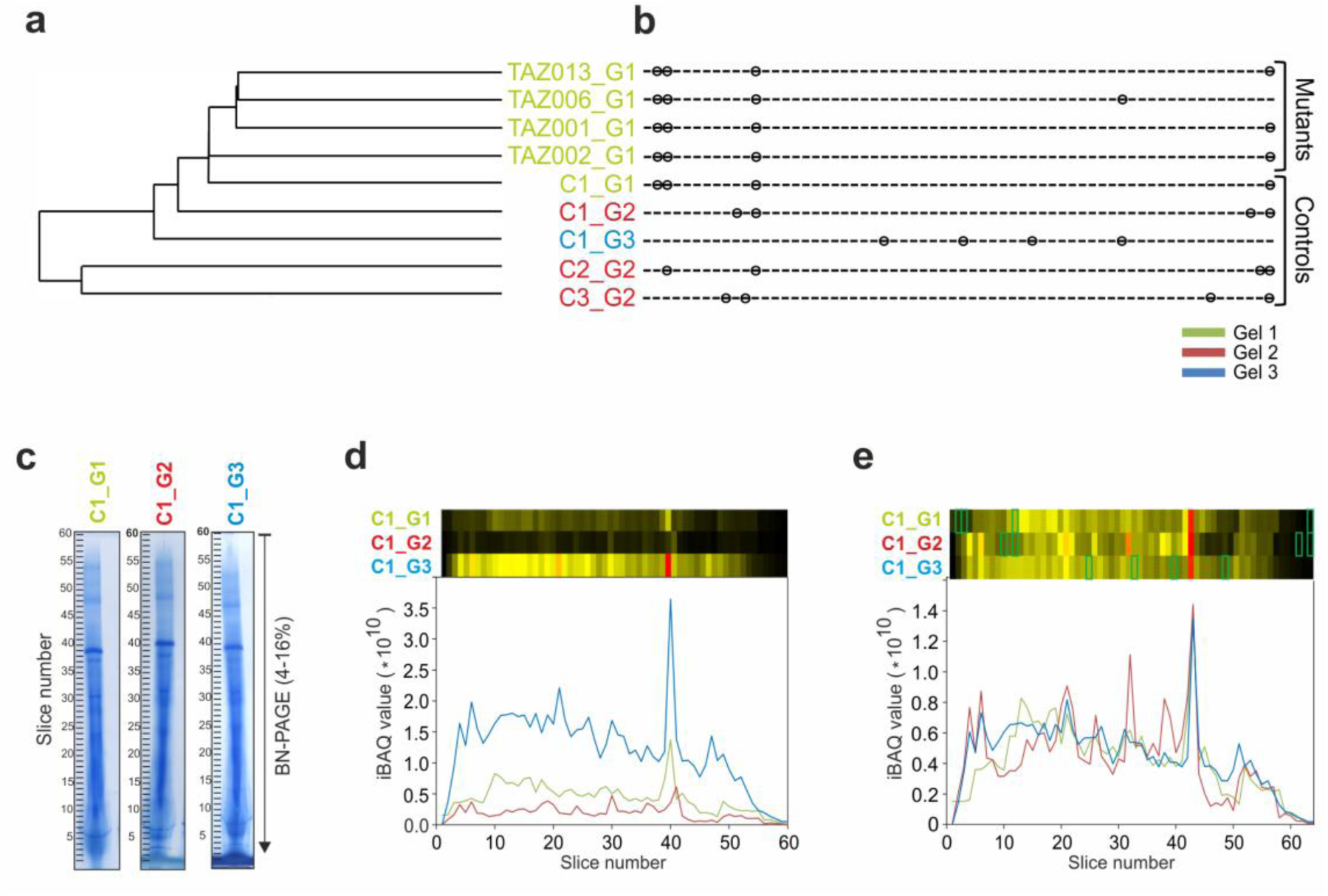
Clustering and alignment of nine complexome profiling examples. The colors refer to the three different BNE gels on which the samples were run. a) single linkage clustering of the profiles based on the total costs obtained from the pairwise alignments. b) Locations of inserted slices after multiple alignment of BTHS mitochondria and control complexome profiles. Complexomes from the same gels tend to have slices inserted at the same locations. c) Images of complexome profiles of mitochondria from the same control on three different BNE gels, illustrating the overall shifts in migration profiles between gels, e.g. at slice 40, d) Summed up intensities (iBAQ values) of all mitochondrial proteins in the original, unaligned and non-normalized gels. The heatmaps were scaled to the maximum over all the gels e) Summed up intensities of all mitochondrial proteins in normalized and aligned gels. The green boxes indicate the positions of the slices added in the alignment procedure.

We first examined how the loss of CL affects the formation of oxidative phosphorylation supercomplexes in the aligned gels, as supercomplexes have been reported to be stabilized by CL (Pfeiffer et al., 2003). In both control and patient samples all complex I was found to be associated with complexes III and IV to form mitochondrial supercomplexes (Schägger and Pfeiffer, 2000) (Figure 3a), which is characteristic for human mitochondria (Acin-Perez et al., 2004; Schägger et al., 2004). The total amount of supercomplexes was hardly changed, but the abundance of higher order supercomplexes containing more than one copy of complex IV (S_2_-S_4_) appeared reduced in mitochondria from patient fibroblasts (Figure 3a). A corresponding increase of complex III dimer at ~500 kDa and III_2_-IV supercomplex at ~700 kDa was observed (Figure 3a). This is consistent with the reported stabilization of respiratory supercomplexes by CL and suggests that the lipid may primarily be important for the interaction of complex IV with the other complexes. It is important to note that the rather subtle decrease in the high-mass shoulder of the supercomplex peak, was retained after alignment and normalization with COPAL and averaging the resulting migration profiles.

**Figure 3.**
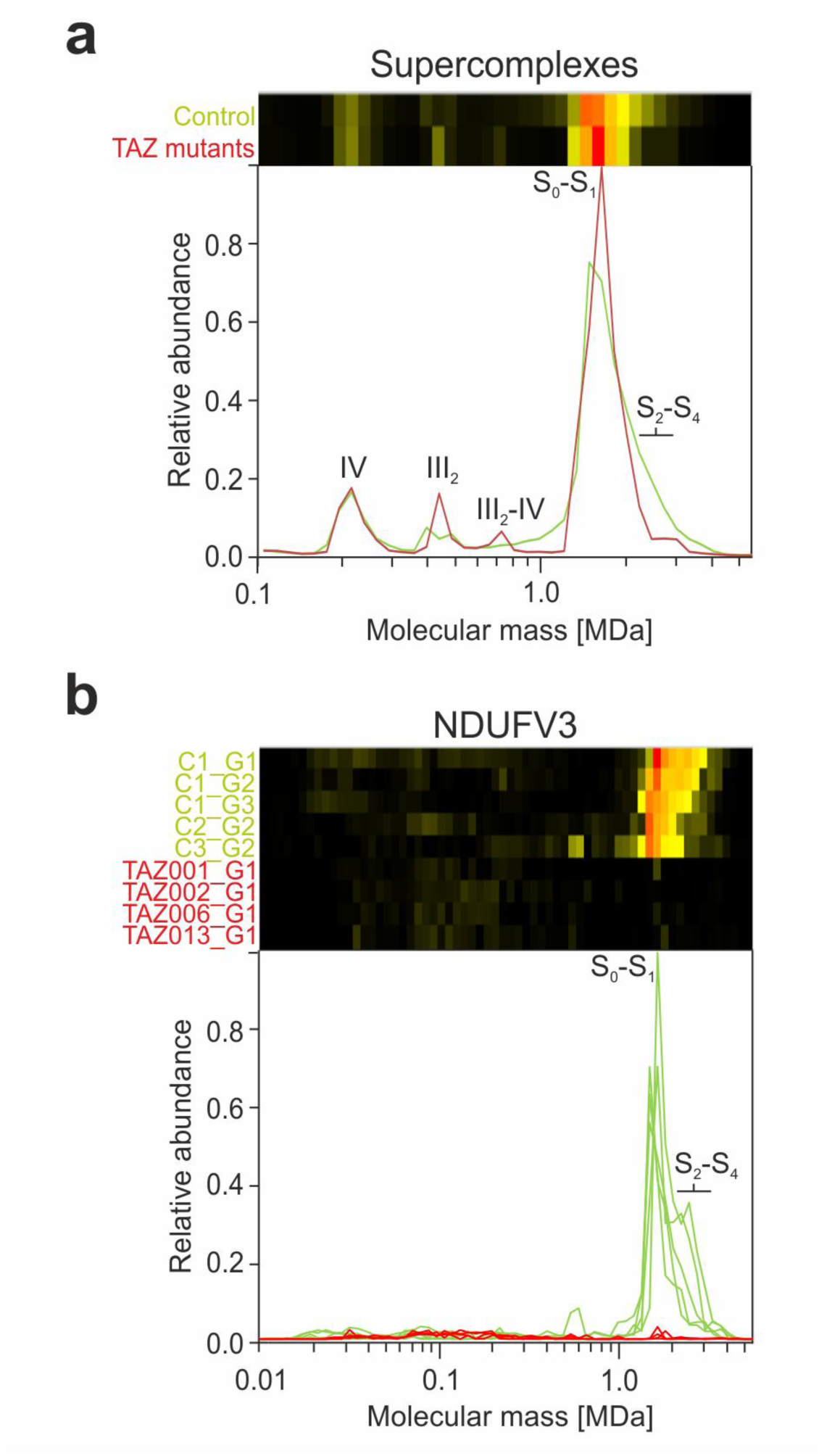
Migration pattern of Oxidative Phosphorylation proteins for controls (green) and TAZ mutation cell lines (red). **a**) The summed intensities of all detected subunits of complexes I, III and IV show that the respiratory chain, including complex I that is part of the S_0-1_ peak, remains intact in the BTHS mitochondria. Nevertheless, changes in the abundance of the complex III dimer, and S_2_-S_4_ supercomplexes are evident when comparing average profiles between BTHS mitochondria and controls. **b**) Levels of the large isoform of complex I subunit NDUFV3 were severely reduced in all BTHS mitochondria profiles, with a small peak still visible at around ~1500 kDa, which corresponds to the peak of supercomplex S_0_-S_1_. Note that there is a substantial amount of variation in the controls for NDUFV3 in the height of the high-mass shoulder diagnostic for supercomplexes S_2_-S_4_.

### Quantifying differences per protein in BTHS mitochondria

To systematically and automatically detect changes common to all variants of *TAZ* after alignment, the complexome profiles were separated into two groups, one containing the four BTHS mitochondria and the other containing the five controls. Hausdorff distances for each protein were calculated for all pairwise comparisons and then used to determine the Hausdorff distance effect size (Methods). The 845 mitochondrial proteins that were detected show a wide range of effects of the mutation (Table S4). Interestingly, one of the highest scoring proteins (rank 3) is the large isoform of NDUFV3 of complex I (NADH:ubiquinone oxidoreductase). It is almost completely absent from the BTHS mitochondria (Figure 3b), even though the assembly and the abundance of complex I was hardly affected (Figure 3a). That NDUFV3 is bound to the part of complex I most distant from the membrane domain with no apparent involvement in supercomplex formation (Guerrero-Castillo et al., 2017b) suggests that the loss of this subunit is linked to the *TAZ* mutations in an indirect way.

### Gene Set Enrichment Analysis

In principle one can combine signals of individual proteins of a protein complex to obtain the average behavior of the complex (Figure 3a). However, this may lead to incorrect quantitative representation of some proteins in the averaged profiles. iBAQ values depend strongly on how well a protein can be detected by mass spectrometry and thus vary greatly between different proteins. Even if the abundance values are normalized between proteins, proteins that occur in different complexes at the same time will become underrepresented in a given multiprotein complex. Therefore, quantitative analyses are commonly done at the level of individual proteins rather than on the averages of proteins within complexes. To obtain statistics on changes at the level of known multiprotein complexes we performed Gene Set Enrichment Analysis (Subramanian et al., 2005) on the list of mitochondrial proteins that were rank ordered based on the Hausdorff effect size. In total five protein complexes had many proteins that scored high on the list and were thus significantly affected in BTHS mitochondria at a P value < 0.001 and a false discovery rate (FDR) < 0.05 (Table 1). Of these affected complexes we will discuss the MICOS and MIB complex in detail.

**Table 1.** Significantly affected protein complexes in BTHS Mitochondria. Proteins were ranked on differences in abundance profiles between BTHS mitochondria and controls, followed by Gene Set Enrichment Analysis (software.boardinstitute.org/gsea/index.jsp). 1000 permutations were performed to obtain normalized enrichment scores and statistical significance values. ES = enrichment score, NES = normalized enrichment score, NOM p-val = nominal P value, FDR q-val = false discovery rate: i.e. after correcting for multiple testing.

| Complex Name | Proteins | ES | NES | NOM p-val | FDR q-val |
| --- | --- | --- | --- | --- | --- |
| 39S ribosomal subunit | 50 | 0,53 | 2,4 | < 0,001 | < 0,001 |
| MICOS complex | 7 | 0,87 | 2,3 | < 0,001 | < 0,001 |
| MIB complex | 12 | 0,74 | 2,3 | <0.001 | <0.001 |
| Prohibitin complex | 2 | 0,99 | 1,7 | < 0,001 | 0.037 |
| 2-oxoglutarate dehydrogenase | 4 | 0,80 | 1,7 | 0,010 | 0.039 |

### The MICOS complex was markedly increased and migrated at a lower molecular mass in BTHS mitochondria

Both the ~700 kDa MICOS complex of the inner mitochondrial membrane and the larger ~ 2 MDa MIB complex in which MICOS is connected to components of the outer membrane (Huynen et al., 2016; Pfanner et al., 2014) were significantly affected in BTHS mitochondria (Table 1, Figure 4a). Separately analyzing the “non-MICOS” constituents of the MIB complex (DNAJC11, SAM50 and MTX1,2,3) revealed that changes in these proteins were barely significant on their own (P=0.055). The significant scores for the MIB complex thus mainly resulted from changes in the components of the MICOS complex. Migration patterns of all seven “classic” components of the human MICOS complex (Pfanner et al., 2014) showed in addition to the expected maxima at about 700 kDa and 2.2 MDa a third peak at ~4.5 MDa most likely corresponding to a dimer of the MIB complex (Figure 4b, c, d). The abundances of the proteins in all three complexes was increased 2 to 6-fold in mitochondria from BTHS fibroblasts as compared to control fibroblasts, except for MIC25 that has a low rank among proteins ordered based on the Hausdorff effect size (Figure 4a). Closer inspection of the aligned and averaged migration profiles revealed further that the maximum of the MICOS complex was downshifted by about 100 kDa to ~600 kDa in the BTHS mitochondria (Figure 4c,d) suggesting that at least one protein was lacking from the ~700 kDa MICOS complex. Three proteins were reported to interact with MICOS: DISC1 (Pinero-Martos et al., 2016), CHCHD10 (Genin et al., 2016) and OPA1 (Glytsou et al., 2016). We detected OPA1 in the fibroblast complexome profiles. Notably, the differences in the migration profiles of OPA1 were among the strongest observed in the dataset (rank 19, Table S4) changing from a broad distribution covering the entire range from ~100 kDa to 4 MDa in the controls to a pronounced single peak in the BTHS mitochondria centered around 100 kDa, matching the mass of the mature monomeric protein (Figure 4d). The disappearance of OPA1 from higher molecular mass complexes may thus provide a straightforward explanation for the 100 kDa lower mass of the MICOS complex in cells with *TAZ* mutations.

**Figure 4.**
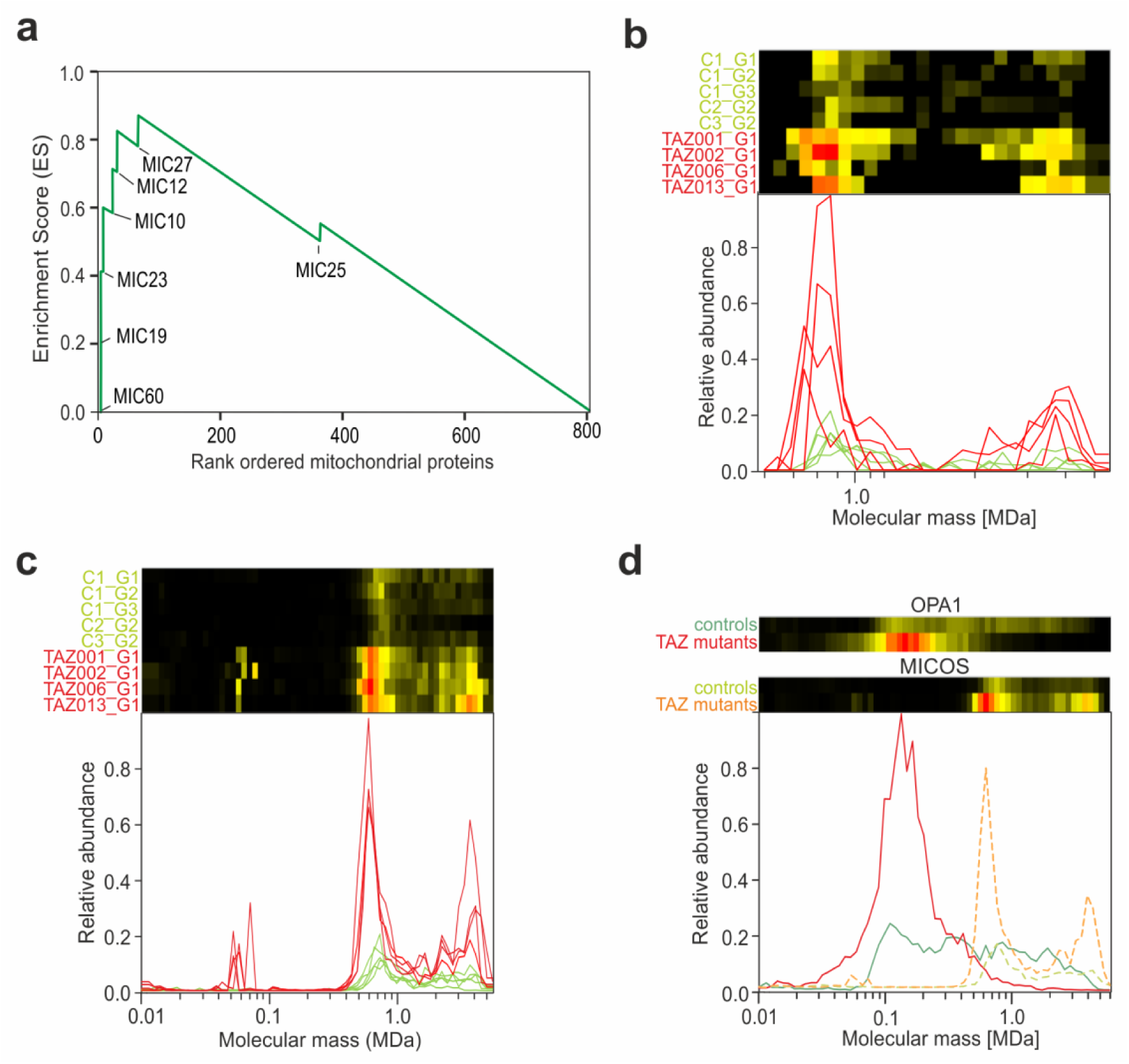
TAZ mutations significantly affect the MICOS complex. a) Gene Set Enrichment Analysis shows that all MICOS complex proteins, except MIC25 that has position 381, rank high on the list of mitochondrial proteins most affected by the *TAZ* mutations and are part of the so-called leading-edge subset in GSEA (Subramanian et al., 2005). b and c) Aligned migration profiles of the MICOS proteins MIC60 and MIC19 respectively. The *TAZ* mutations lead to an increase in the expression of both proteins and to a small shift to a lower molecular mass of the complete complex from ~700 kDa to ~600 kDa. For MIC19 they lead to a new peak around a molecular mass of 58 kDa, potentially reflecting a new subcomplex, possibly a MIC19 dimer as the predicted molecular mass of mature MIC19 is 23 kDa. d) Average migration profile of all MICOS proteins and OPA1 in controls and in BTHS mitochondria. The BTHS mitochondria display, relative to controls, a higher expression of MICOS proteins and a shift to a lower mass of the MICOS complex. OPA1 changes from a broad distribution in the controls to a pronounced single peak in the BTHS mitochondria centered around 100 kDa.

## DISCUSSION

The comparison of complexome profiles holds the promise to systematically map how metabolic or genetic challenges affect the composition, abundance and stability of protein complexes and whether there are tissue specific variations in protein complex compositions. To this end, we implemented a combination of bioinformatic approaches in COPAL, allowing large scale comparative analyses of complexome profiles. It is instructive to examine the differences between the results obtained with COPAL and the subsequent gene set enrichment analysis with those obtained using expert, manual curation. In a previous study, the complexome profiles obtained from BTHS mitochondria were manually compared with that of a control that was run on the same gel (C1_G1) (Chatzispyrou et al., 2018). As expected, most of the prominent changes were detected by both approaches, however the example of the MICOS complex (Rampelt et al., 2017) showed that application of COPAL to quantitatively compare multiple profiles allows a more detailed analysis revealing features not evident from manual inspection. While the overall increase in abundance of the canonical MICOS proteins was easily observed by visually comparing the individual profiles (Chatzispyrou et al., 2018), the shift in its size amounting to only about 15% of its total mass and allowing us to identify loss of OPA1 as likely candidate explaining this mass difference was only detected following alignment of multiple proteins by COPAL. Notably, the shift of two gel slices (~100 kDa) is smaller than the shifts that COPAL introduces to align the complexome profiles (four gel slices), underlining the relevance of such alignment to detect changes in molecular mass.

Aside from the change in the MICOS complex abundance also the 2-oxoglutarate dehydrogenase complex (Figure S2) and prohibitin (Figure S3) were detected in both analyses. The decrease in higher order respiratory chain supercomplexes caused by the mutations was not detected in a global automated analysis and probably would have been missed by manual inspection as well without knowing from earlier studies that CL is involved in the supercomplex assembly and stability (Mileykovskaya and Dowhan, 2014; Pfeiffer et al., 2003). On the other hand, the fact that the decrease in the high-mass shoulder of the major supercomplex peak was still visible following application of COPAL and averaging of the migration profiles demonstrated the feasibility the alignment. The complete disappearance of a ~ 3 MDa branched-chain amino acid dehydrogenase complex reported by Chatzispyrou *et al*. (Chatzispyrou et al., 2018) was not detected because there was a large variation in the presence of the ~3 MDa (DBT and BCKDK) assembled complex among the controls and because the dominant peak for other proteins remained at the same mass (BCKDHA, BCKDHB) (Figure S4). Conversely COPAL did detect significant differences in the large subunit of the ribosome that was not uncovered in the manual examination (Figure S3). This appears mainly the result of COPALs ability to automatically and unbiasedly process and prioritize large numbers of proteins.

Our study provides proof-of-principle for the usefulness of COPAL to analyze multiple complexome profiling datasets. It shows that COPAL can be a useful tool to combine and compare biological or technical repeats of all kinds of complexome profile experiments. Thereby, it will allow the large scale comparative analysis of complexome profiles, facilitating robust assessment of complexome inventories and allowing detailed and reliable conclusions.

## Supplementary material

Supplementary tables with the original (Table S2) and aligned (Table S3) complexome data are available from the corresponding author.

## Acknowledgements

This work was supported by a TOP grant from the Netherlands Organization for Health Research and Development (no. 91217009) to M.H. and U.B.

